# Population estimation study reveals staggeringly high number of cattle on the streets of urban Raipur, Chhattisgarh, India

**DOI:** 10.1101/2020.06.01.127506

**Authors:** Bhupendra Kumar Sahu, Arti Parganiha, Atanu Kumar Pati

## Abstract

Cattle (bovine species) are economically and ecologically very important and are cosmopolitan in distribution. Increasing number of cattle on the urban streets of developing and underdeveloped countries has become an unmanageable menace in recent time. Consequently numbers of road accidents have increased in the urban areas of most of the developing countries, like India. In the present study, we estimated the population of street cattle wandering on the street/road/highway of Raipur city of India using direct head count method and advanced Photographic Capture Recapture Method (PCRCM). We compared these two scientific methods of population estimation to check their adequacy. We prepared grid (1.0 x 1.0 km) on the map of Raipur city using Quantum Geographic Information System (QGIS) software and randomly selected 20 grids for the estimation of street cattle population. We used line transects and block count methods for data sampling. Results of direct head count method indicated an estimation of 11808.45 cattle (using line transects sampling method) and 11198.30 cattle (using block counting sampling method) visibly roaming on the street of Raipur city. Further, advanced PCRCM indicated an estimation of 35149.61 cattle using line transects sampling method and 34623.20 cattle using block counting sampling method. We observed female biased sex ratio in both mature and immature cattle. Frequency of mature cattle was significantly higher than that of immature cattle followed by calves. Further, the frequency of cattle in a grid was found in the following order: cow > bull > heifer > immature male > female calve > male calve. We concluded that the estimated population of street cattle in Raipur city is about 34623. Results of both the techniques, i.e., direct head count method and PCRCM for population estimation are consistent. The direct head count method yields the number of cattle visibly roaming on the street in a particular time; whereas advanced PCRCM gives the total population of street cattle in the city. Results of this study might be helpful in the management of street cattle menace in urban habitat and landscape.

## Introduction

Man began to domesticate cattle about 10000-to 8000-year ago [1,2]. They used them as the sources of milk and meat. Cattle are distributed worldwide and constitute the largest number among the ungulates [3,4]. In the animal world, cattle form the maximum zoomass, i.e., 600 million metric tons [5].

At the end of 2018, worldwide cattle population was about 996.36 million [3]. As per worldwide cattle inventory 2018, India is the leading country with the largest number of cattle (i.e., 305.00 million) in the world [4]. This figure was about 196.19 million during 2012 (19^th^ livestock census, 2012) that included 5.29 million stray (free roaming ownerless) cattle [6]. Recently, 20^th^ livestock census of India (2019) was released and it showed that cattle population is about 198.48 million that included 5.02 million stray cattle in India [7]. In Chhattisgarh state, cattle population is about 9.98 million including 3.74 lakh in urban areas; and the stray cattle is about 1.85 lakh [7]. As a consequence of increasing cattle population, one can find a large number of cattle wandering here and there on streets in most of the developing and underdeveloped countries [8–11]. Increased number of street cattle could be attributed to the disappearance of grassland on account of rapid urbanization of land area by both the government and private sectors. Due to shortage of the foraging spots, cattle wander freely on the streets of urban areas. One of the important reasons of increased number of cattle on urban street is that due to space crunch people do not prefer to keep their domestic cattle at the home. Owners release their cattle on the nearby streets either temporarily or permanently.

People in urban areas share public places with bovine species. Both of them interact directly or indirectly and face a lot of problems [8–11] Due to lack of natural diet, cattle consume harmful materials, like polythene, heavy metals, and so on [12–14]. Cattle can neither digest nor egest polythene/plastics materials. These harmful materials stuck into their stomach that eventually causes cancer and leads to early death. About 95% to 100% of street cattle ingest these hazardous materials into their stomach [12,13]. These harmful materials and their metabolites slowly sip in to the human system through milk and milk products leading to several health-related consequences. Presence of cattle on the street cause increased road accidents in urban areas. People do not feel safe when walking or riding two/four wheeler on the road. It has been reported that about 300 people died during the last 2.5 years in the State Punjab, India on account of stray cattle menace on the road [15].

Management of street cattle is one of the very important issues in many countries, including India. Management of these cattle can be effective only when we know adequately about the population of street cattle and their behavioural disposition.

Population estimation of cattle in India is mostly based on door to door inventory collected from the owners of the cattle. This may not provide the accurate number of cattle present on the street especially in urban areas where people do not like to keep their cattle at home and try to mislead investigating officers through providing false information.

There are several methods to estimate the population of mammalian species. Capture Mark Recapture Method (CMRCM) is very popular and is a widely accepted technique. In this technique, animals are first captured and marked with either color [16–19] or some tags attached to the animals [20]. After a desirable time lag (in days), animals of the same area are recaptured. On the basis of number of animals on two different days with the number of previously marked animals on recapture day, the population of animals is estimated. The estimation by the Lincoln index is considered useful in this context. Lincoln index is one of the widely used and accepted indices in ecology to estimate the population of any species. The color or physical tags used in the CMRCM for identifying individuals may alter the physiology and behaviour of the studied animals [20] and sometimes even lead to death of the animals. This fatality could be attributed to hazardous nature of the colored chemical and its amount used for tagging. The other negative aspect of this method is that animals become vulnerable to their predators because of the marker elements.

In the advanced technique of CMRCM known as Photographic Capture-Recapture Method (PCRCM), marking animals with the hazardous colored chemicals or tag is not required. In this technique, the natural color pattern [21,22] or other biographical and morphological characteristics [9,23,24] of the animals are used to identify the individuals without any direct/indirect harmful impact on them.

Another method used in recent studies is the direct head count method [25,26] to estimate density as number of heads per kilometer distance. It is very similar to the line transect sampling method [27]. Head count sampling can be carried out using either block count sampling method or line transect sampling method [27]. Density is determined by these sampling methods. One can estimate the total population by multiplying the density to the total area or distance/length of the routes [25,27].

In the current study, we estimated the population of street cattle wandering on the street/road/highway of Raipur city (primary objective) and validated the appropriateness of the direct head count method by comparing our results with the results of well established PCRCM (secondary objective).

## Material and methods

### Study area and subject

Study was carried out in Raipur city (Latitude: 21.2514° N; Longitude: 81.6296° E, and Elevation: 298 m from the sea level). It is the largest city and the capital of Chhattisgarh (a state of the Indian Federal Union). The total length of street/road/highway of Raipur city is 1622.04 km distance and transect over 226 km^2^ area of the city as per Municipal Corporation Raipur [28]. The human population of the city in 2011 was 1.01 million [29]. Now it has increased to about 1.87 million in 2019 [30].

Cattle wandering, foraging, standing or laying on the street of the city of Raipur were used as the subjects in the current study. We considered all the cattle of different age classes as street cattle that were seen on the street/road/highway and within adjacent 10 meters on both sides during our observation. These cattle are either stray cattle or free roaming owned cattle released in the road permanently or temporary basis.

### Preparation and selection of grid

We procured the map of Raipur city from the Municipal Corporation Raipur. Google map of Raipur city was also accessed to find out route of highway/road/street. We prepared grid (1.0 x 1.0 km) on Google map of Raipur city (Fig 1) using Quantum Geographic Information System (QGIS) software [31]. The total grids counted those overlapped the map of Raipur city were 190. In this study, we included the grids those covered ≥ 50% of area (i.e., 163 grids: 151 grids covered 75-100% and 12 grids covered 50-75%). we excluded remaining 27 grids (i.e., the grids those covered < 50% of area). Further, each grid was provided with a Grid ID number from 1 to 163. Twenty out of 163 grids were randomly selected for the sampling of street cattle population. We generated 20 random numbers from among 1 to 163 by random number generation technique using MS Excel.

**Fig 1.**
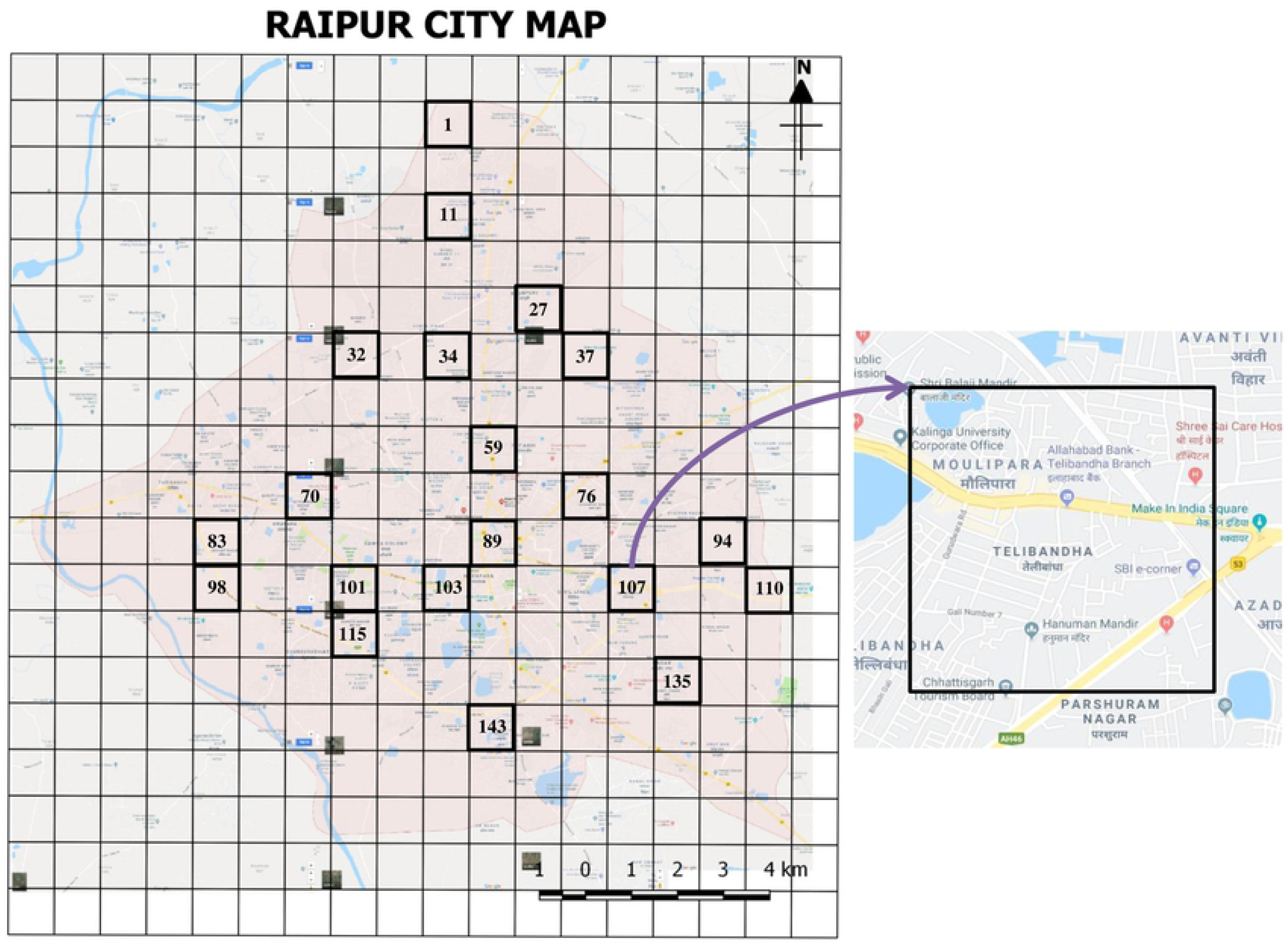
Study location and grid. The picture illustrates layering of 1.0 km^2^ grids over a static Google map of the Raipur city. Although 190 grids overlap areas of the city on the map, only 163 grids were selected for the study. Twenty-seven (27) grids were excluded as they were covering less than 50% grid area on the map. However, grids covering more than 50% grid area on the map were included. Of the 163 grids, only 20 grids were chosen randomly for the study. The selected grids were numbered (Grid ID #1, 11, 27, 32, 34, 37, 59, 70, 76, 83, 89, 94, 98, 101, 103, 107, 110, 115, 135, and 143). The Grid #107 has been shown in the inset with road map within the grid area.

### Sampling method

We carried out the study between February 16, 2019 and April 11, 2019. We followed line transect [25,27] and block count methods [27] for sampling of the data. During sampling hours, direct observation of cattle was made by two observers. We followed all routes of highway/road/street of the selected grid using the Google map and Map Ruler Lite (https://apkpure.com/maps-distance-ruler-lite/com.mobilerise.MapsRuler2) [32] mobile apps by walking and driving/riding a motorcycle. We sampled data at two different days. On the first day, we photographed the street cattle using Nikon D-3400 still camera at the selected grid covering all the routes and on the third day we took photographs again (recaptured) at the same grid approximately at the same time. We repeated the same protocol for all 20 selected grids. We started and ended the photography of the street cattle from the predefined starting point to end point of the sampling grid, respectively. We followed the same route of grid in unidirectional way during both capturing and recapturing days. We ensured to examine each route but avoided repeated observation of the same route and double count of the cattle. Biographical (gender, age class), morphological (body color, horn shape & size, udder size), and additional characteristics (collar, bells and cut mark in ear) of each cattle were also noted down during sampling (Fig 2 and Table 1).

**Table 1.** Biographical, morphological, and additional characteristics/traits used for identification of individual street cattle in Raipur city using advanced Photographic Capture-Recapture Method (PCRCM).

| Characteristics/traits | Categories |
| --- | --- |
| Biographical |  |
| Gender | Male/Female |
| Age class | Male: Bull/Immature male/Male calve<br>Female: Cow/Heifer/Female calve |
| Morphological |  |
| Color | Main body color<br>Patch/ mark color |
| Udder size of female | Largest/Larger/Large/Medium/Small/Smaller/Smallest |
| Presence of horn | Present/Absent |
| Size of horn | Largest/Larger/Large/Medium/Small/Smaller/Smallest |
| Direction of horn | Upward/Downward/Forward/Backward/Sided |
| Broken horn | Left horn/Right horn |
| Additional characteristics |  |
| Presence of collar/rope | Present/Absent<br>If present, Color of collar/rope |
| Presence of bell/rope in collar | Present/Absent |
| Presence of ear-tag or cut-mark in ear | Present/Absent |

**Fig 2.**
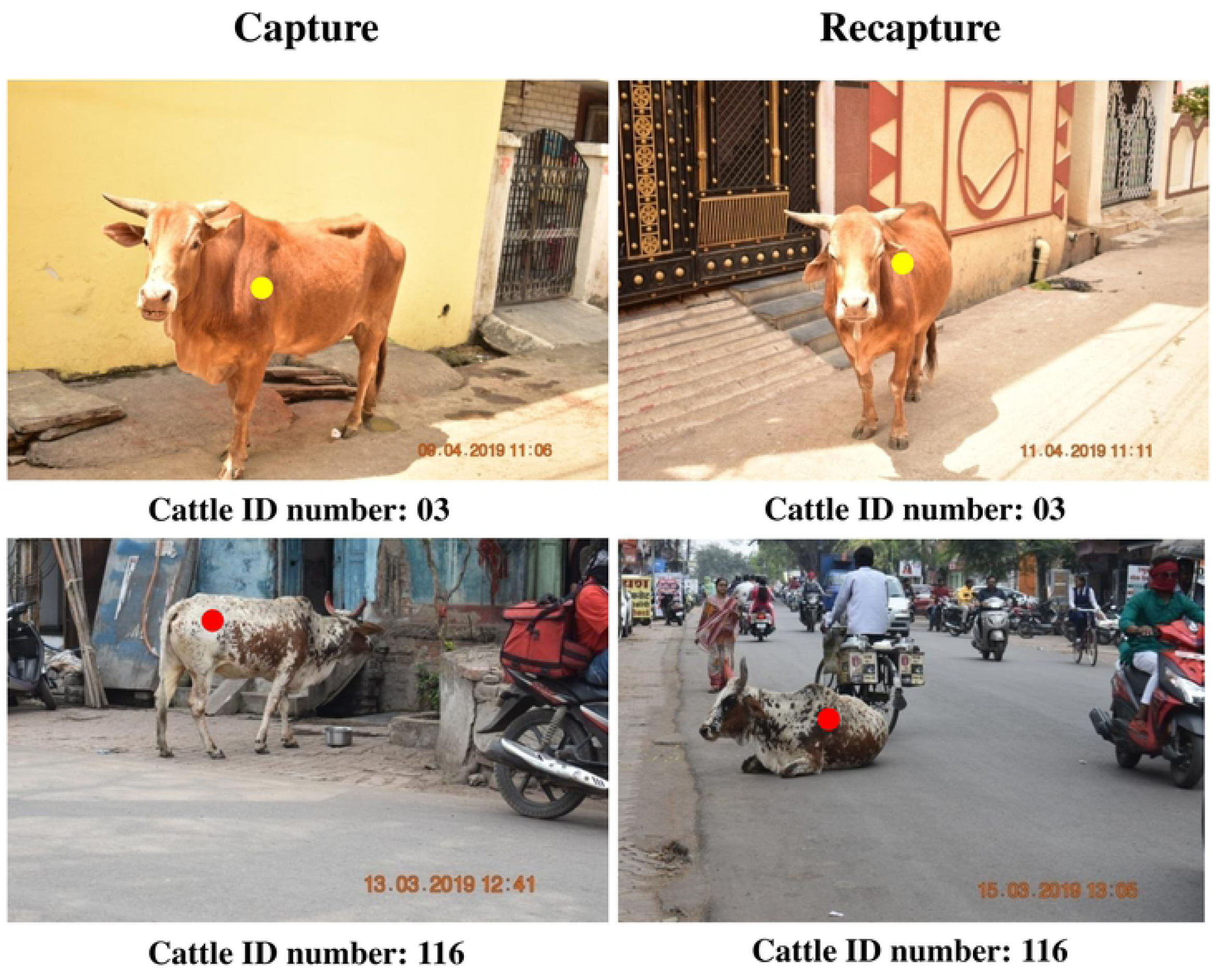
Photography and identification of street cattle. An illustrative example of street cattle showing two different street cattle (Cattle ID #03 and #116) that were photographically captured on the first day and recaptured on the third day of the sampling in their respective grids. Identification was done by their biographical (gender, age class), morphological (body color, horn shape & size, udder size), and additional characteristics (collar, bells and cut mark in ear).

We carried out our field work between 1100 h and 1500 h to increase the sightability of the subject [22]. At this time point of the day, mostly street cattle are active and busy in foraging on roadside open garbage and dumping bins [9,11].

### Ethical approval statement

Ethical approval was not required in this study as we only photographed the animal subject. Our study protocol did not involve handling of animal for any purpose. We did not harm the cattle directly or indirectly for any rationale of the study. During photography, we always maintained 10 m distance from the animal subject so that even their behaviour could not be disturbed. The study was carried out in public places within the city, therefore, we obtained permission from the Municipal Corporation, Raipur, Chhattisgarh [Ref. number: 170/health department/MCR/2018; dated: 15/03/2018].

## Statistical analysis

### Street cattle population size estimation

To estimate the population of street cattle we used direct head count method and Photographic Capture-Recapture Method (PCRCM). In PCRCM, for the calculation of size of the population, we excluded the data of Jai Stambh Area (Grid - 89) because the number of street cattle was undefined as per the Lincoln index formula when common street cattle in two sampling days is found to be zero. However, data of this area was used in the calculation of direct head count method.

#### Method 1: Direct head count method

We observed street cattle visibly roaming on street using two sampling methods. First, line transect sampling method [25,27] and the second block count sampling method [27]. To estimate the total population we first estimated the density of street cattle.

Density of cattle can be defined as outlined below:

Number of cattle per kilometer distance (for line transect sampling method):

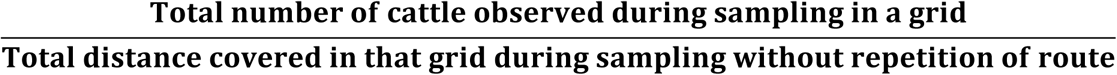

Subsequently we estimated the *total number of cattle visibly roaming on the street* of Raipur city using the formula:

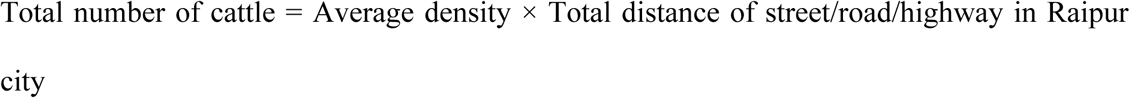

Number of cattle per square kilometer area (for block count sampling method):

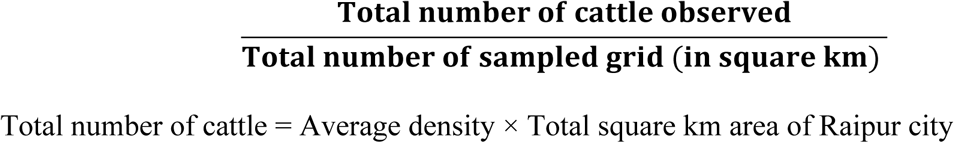

#### Method 2: PCRCM method & Lincoln index formula

Photographic Capture-Recapture Method (PCRCM) and Lincoln index formula were mutually used to estimate the population of street cattle in 19 studied grids. We first estimated the population of street cattle in 19 different grids using the Lincoln index formula:

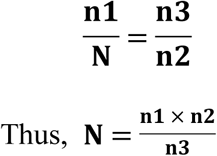

Where,

n1 = Number of cattle photographed on the first day;

n2 = Number of cattle photographed on the third day;

n3 = Number of cattle sighted on the first day and also on the third day;

N = Estimated population of cattle in a grid (measured in 1km^2^ area)

We also estimated average number of street cattle in unit square kilometer area (average density). Thereafter, we estimated the *total population of street cattle in Raipur city* using the following formula:

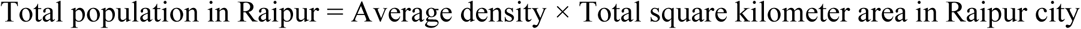

#### Other population ecological variables

We further analyzed gender composition, sex ratio, and age composition of the street cattle.

We employed one way ANOVA to compare the density of cattle “cattle per km” in 20 different studied grids. We performed Mann-Whitney U test to compare frequency of male and female cattle. We set the significant level at *p* ≤ 0.05. Sex ratio was derived from the number of male street cattle divided by number of female street cattle.

Normality and homogeneity of the data were checked using Kolmogorov-Smirnov statistic and Levene Statistic, respectively. Non-parametric ANOVA (Kruskal-Wallis rank test) followed by Mann-Whitney U test was performed to compare the frequency of cattle at three different age groups (mature, immature, and calve) and frequency of six different cattle groups (cow, bull, heifer, immature male, female calves, and male calves). Bonferani adjustment was used in both the cases. We analyzed the data with the help of SPSS (version 20.0).

## Results

### Population density of cattle

#### Method 1: Direct head count method

Table 2 represents the results of direct head count method. The total head counts during the first and the third day were 998 and 984, respectively. Average number of street cattle was 991 and observed average density in 20 studied grids was 49.55 cattle km^-2^ area as per block count sampling method. Further, we covered a total of 141.68 km in all 20 grids during capture on the first day, and repeated the same route during recapturing on the third day. According to line transect sampling method the observed average number and average density of street cattle in all 20 grids were 145.53 and 7.28 km^-1^ distance, respectively.

**Table 2.**
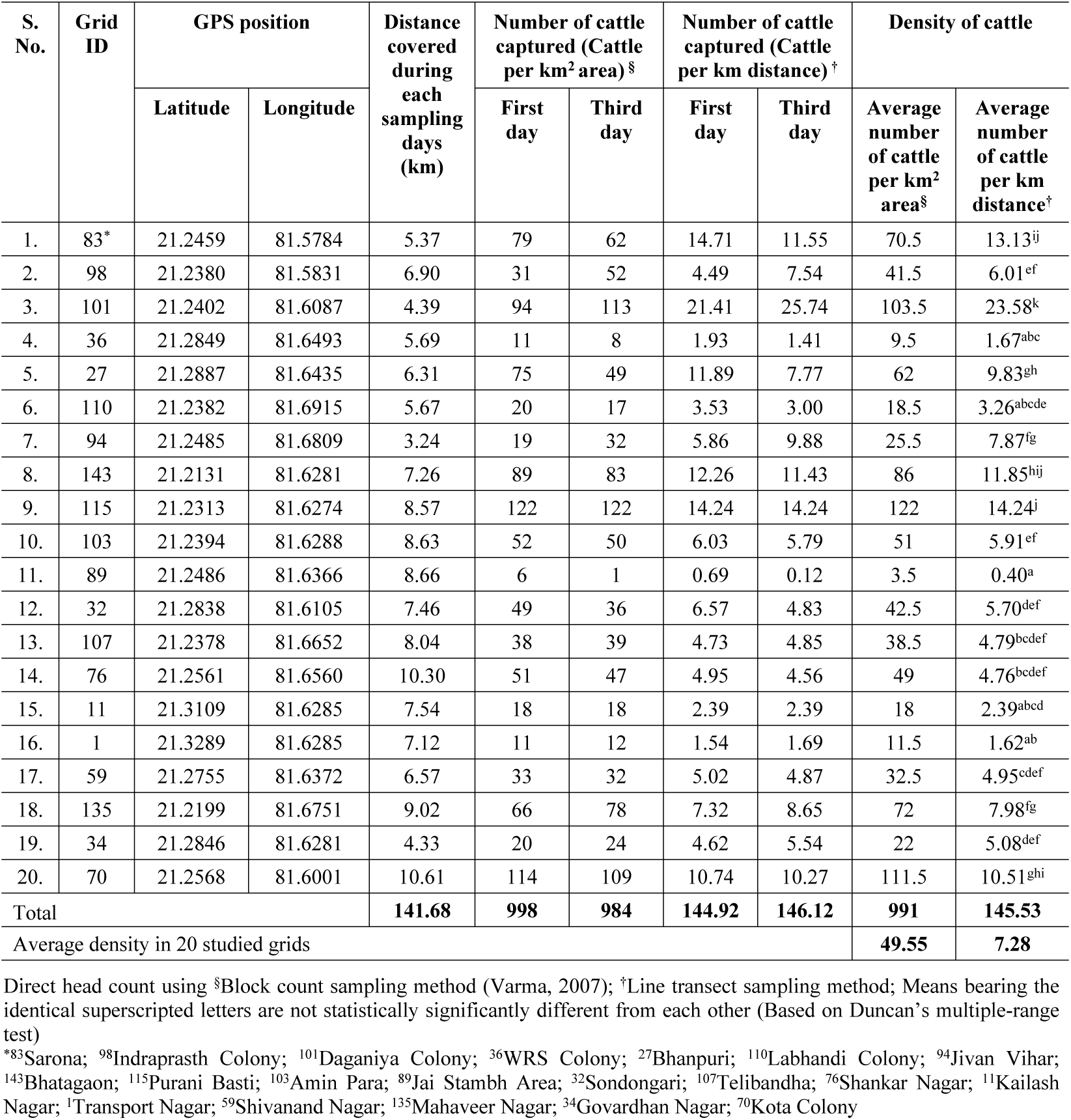
Density of street cattle: average number of cattle per km^2^ area^§^ and per km distance^†^ in Raipur city based studies conducted at 20 randomly selected grids.

#### Method 2: PCRCM method & Lincoln index formula

We estimated the population of street cattle in 19 studied grids separately using the advanced PCRCM and Lincoln index (Table 3). The total estimated population in all 19 grids was 3195.92 street cattle in per km^2^ area. In the unbiased estimation, the estimated population was 2910.85 street cattle in 19 studied grids. The maximum estimated population, i.e., 367.50 street cattle was found in Grid - 27 (Bhanpuri), and the minimum population, i.e., 14.67 street cattle was estimated in Grid - 36 (WRS Colony). Further, the estimated average density of street cattle was 168.21 per km^2^ in 19 sampled grids. On the other hand, according to unbiased estimation the estimated average density of street cattle was 153.20 per km^2^.

**Table 3.** Estimated population of street cattle per km square area using advanced Photographic Capture-Recapture Method (PCRCM) & Lincoln index at 20 different sampling grids in Raipur city.

| S. No. | Grid Number | Capturing Day (CD) | Recapturing Day (RCD) | Day 1 (cattle per km<sup>2</sup>) n1 | Day 3 (cattle per km<sup>2</sup>) n2 | Common in day1 and day 3 (cattle per km<sup>2</sup>) n3 | Estimated cattle population in per km<sup>2</sup> area<br>N= n1×n2/n3 |
| --- | --- | --- | --- | --- | --- | --- | --- |
| 1. | 83 | 14/02/2019 | 16/02/2019 | 79 | 62 | 20 | 244.90 |
| 2. | 98 | 18/02/2019 | 20/02/2019 | 31 | 52 | 12 | 134.33 |
| 3. | 101 | 22/02/2019 | 24/02/2019 | 94 | 113 | 39 | 272.36 |
| 4. | 36 | 25/02/2019 | 27/02/2019 | 11 | 8 | 6 | 14.67 |
| 5. | 27 | 25/02/2019 | 27/02/2019 | 75 | 49 | 10 | 367.50 |
| 6. | 110 | 26/02/2019 | 28/02/2019 | 20 | 17 | 6 | 56.67 |
| 7. | 94 | 09/03/2019 | 11/03/2019 | 19 | 32 | 4 | 152.00 |
| 8. | 143 | 12/03/2019 | 14/03/2019 | 89 | 83 | 25 | 295.48 |
| 9. | 115 | 13/03/2019 | 15/03/2019 | 122 | 122 | 52 | 286.23 |
| 10. | 103 | 16/03/2019 | 18/03/2019 | 52 | 50 | 21 | 123.81 |
| 11. | 89* | 19/03/2019 | 21/03/2019 | 6* | 1* | 0* | undefined* |
| 12. | 32 | 20/03/2019 | 22/03/2019 | 49 | 36 | 12 | 147.00 |
| 13. | 107 | 23/03/2019 | 25/03/2019 | 38 | 39 | 20 | 74.10 |
| 14. | 76 | 26/03/2019 | 28/03/2019 | 51 | 47 | 11 | 217.91 |
| 15. | 11 | 27/03/2019 | 29/03/2019 | 18 | 18 | 4 | 81.00 |
| 16. | 1 | 30/03/2019 | 01/04/2019 | 11 | 12 | 2 | 66.00 |
| 17. | 59 | 03/04/2019 | 05/04/2019 | 33 | 32 | 7 | 150.86 |
| 18. | 135 | 04/04/2019 | 06/04/2019 | 66 | 78 | 29 | 177.52 |
| 19. | 34 | 08/04/2019 | 10/04/2019 | 20 | 24 | 6 | 80.00 |
| 20. | 70 | 09/04/2019 | 11/04/2019 | 114 | 109 | 49 | 253.59 |
| Total estimated population in 19 km<sup>2</sup> area |  |  |  |  |  |  | **3195.92** |
| Average density |  |  |  |  |  |  | **168.21** |
| Unbiased estimated population in 19 km<sup>2</sup> area |  |  |  | 992 | 983 | 335 | **2910.85<sup>a</sup>** |
| Unbiased average density |  |  |  |  |  |  | **153.20** |

<sup>n1</sup>Number of cattle photographed on the first day; <sup>n2</sup>Number of cattle photographed on the third day; <sup>n3</sup>Number of cattle sighted on the first day and also present on the third day; <sup>N</sup>Estimated total population of cattle in 19 grid (1.0 x 1.0 km); \*grid number 89 was not included in the calculation because the number of street cattle was undefined as per the Lincoln index formula when common street cattle in two sampling days is found to be zero, <sup>a</sup>unbiased estimation

## Estimated population size of street cattle in Raipur city

### Method 1: Direct head count method

According to the Municipal Corporation, Raipur, there is 1622.04 km street/road/highway network in 226 km^2^ area in Raipur city. By line transect sampling method we estimated 11808.45 streets cattle in Raipur city. By block count sampling method the estimated street cattle in 226 km^2^ area was 11198.30 (Table 4). Here both the estimated population was derived without using Lincoln index.

**Table 4.**
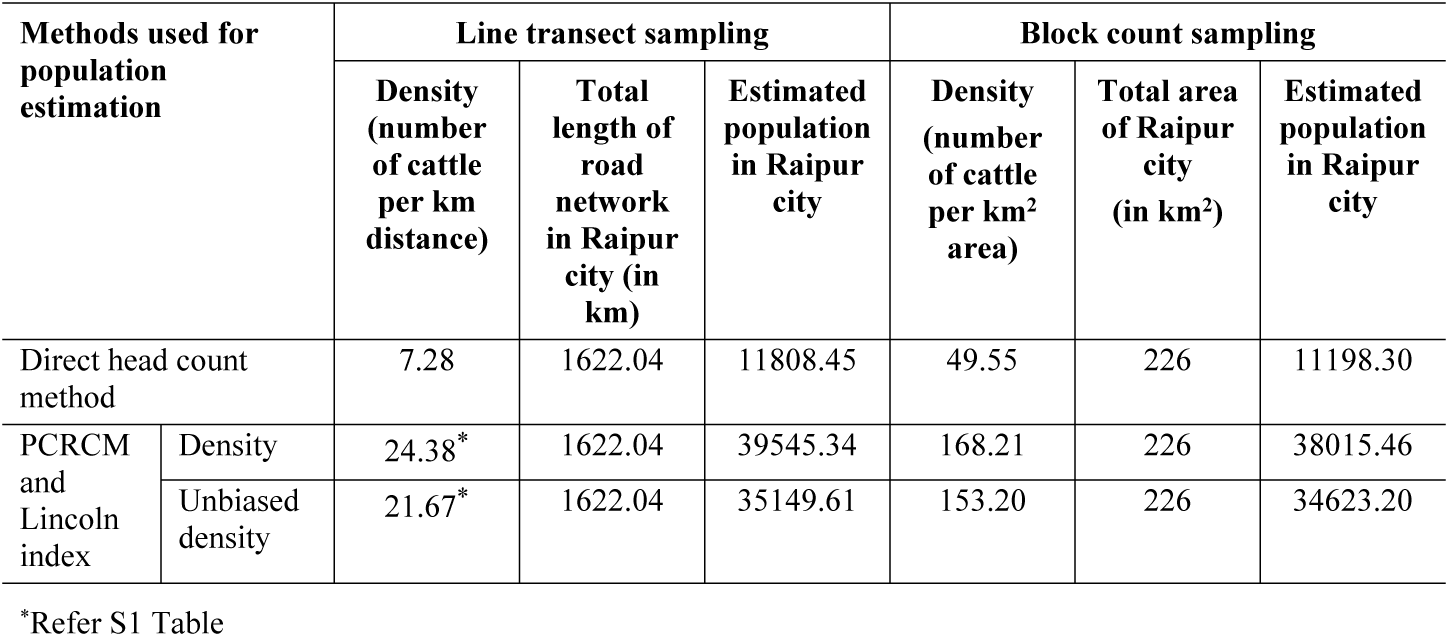
Estimated total population of cattle in Raipur city using two recommended methods, namely Line transect sampling and Block count sampling.

| Methods used for population estimation |  | Line transect sampling |  |  | Block count sampling |  |  |
| --- | --- | --- | --- | --- | --- | --- | --- |
|  |  | Density (number of cattle per km distance) | Total length of road network in Raipur city (in km) | Estimated population in Raipur city | Density (number of cattle per km<sup>2</sup> area) | Total area of Raipur city (in km<sup>2</sup>) | Estimated population in Raipur city |
| Direct head count method |  | 7.28 | 1622.04 | 11808.45 | 49.55 | 226 | 11198.30 |
| PCRCM and Lincoln index | Density | 24.38* | 1622.04 | 39545.34 | 168.21 | 226 | 38015.46 |
|  | Unbiased density | 21.67* | 1622.04 | 35149.61 | 153.20 | 226 | 34623.20 |

\*Refer S1 Table

### Method 2: PCRCM method & Lincoln index formula

The estimated population of street cattle in the total area (226 km^2^) of Raipur city was 38015.46. On the other hand, according to unbiased estimation, the estimated population of street cattle of the city was 34623.20 (Table 4). This result was based on block count sampling method. Table 4 also revealed a comparative output of the direct head count method and PCRCM method. We found that the former method underestimated the population and just showing the cattle visibly roaming on the street, whereas the latter method estimated the population of street cattle in the whole city.

## Results of other population ecological variables

### Effect of the factor “location” on density of street cattle

Cattle per km in different studied grid is significantly different (*F*_19,20_ = 30.24; *p* < 0.001). As per line transect sampling method, the average number of cattle per km distance was found to be the maximum (23.58 cattle per km) at Grid - 101 (Daganiya Colony) and the minimum (0.40 cattle per km) at Grid - 89 (Jai Stambh Area). On the other hand, as per the block count sampling method we found the maximum density (122 cattle per km^2^) at Grid - 115 (Purani Basti) and the minimum density (3.5 cattle per km^2^) at Grid - 89 (Jai Stambh Area) (Table 2).

### Gender composition

Fig 3 depicts the results of Mann-Whitney U test. Results indicate that the overall number of sighted female cattle was significantly higher as compared to that of the male cattle (U = 431.50; *p* < 0.001). We also analyzed sex ratio at different age levels and compared them with each other. The number of male cattle was significantly lower than female cattle in two age groups, i.e., in mature group (U = 379.50; *p* < 0.001) and immature group (U = 508.00; *p* < 0.05). However, this difference was not significant in the calve group (U = 652.00; *p* > 0.05).

**Fig 3.**
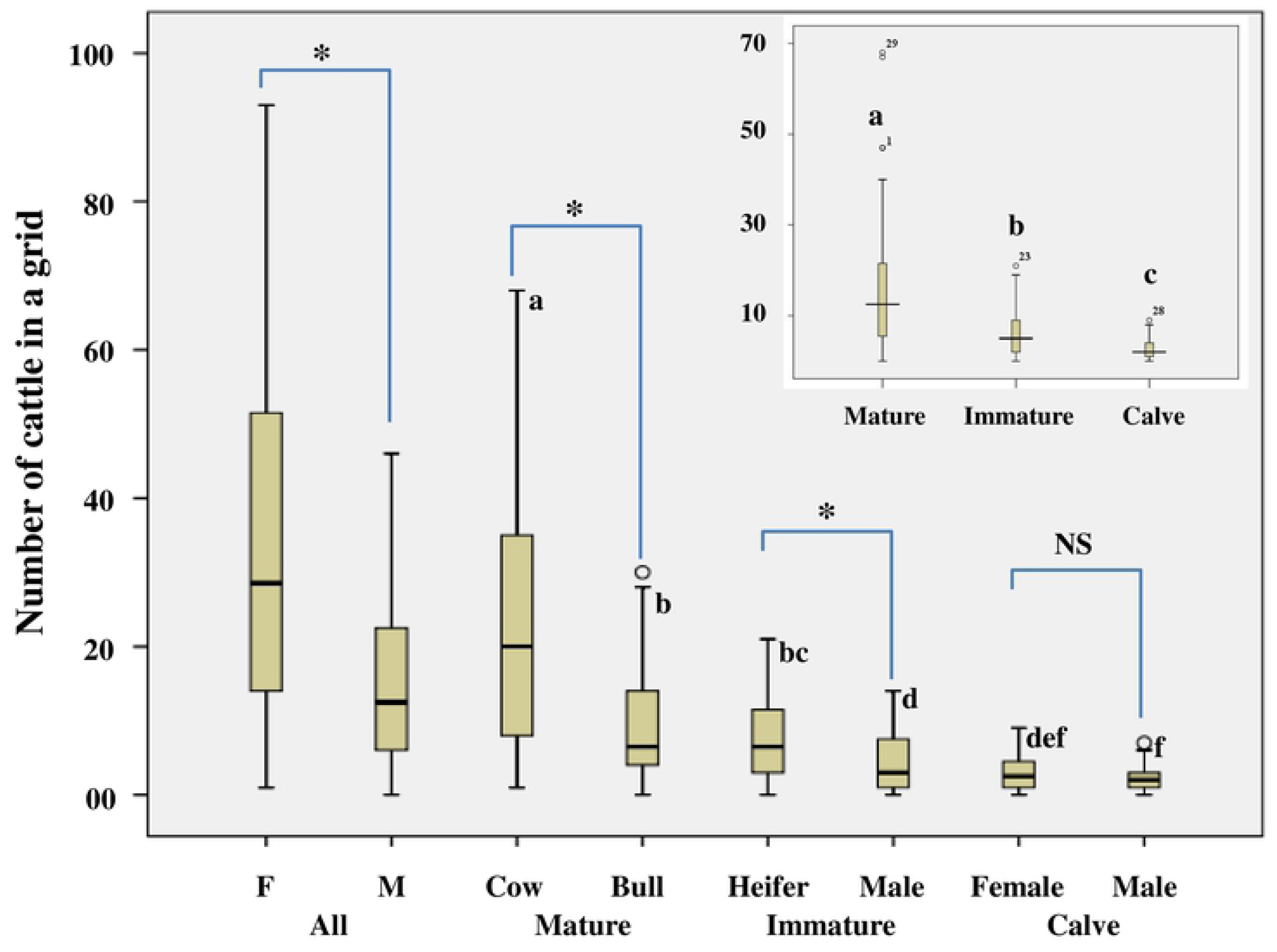
Box plots depicting the number of cattle in a grid by age groups and by gender. The horizontal line that divides the box into two parts represents the median value. The upper and lower limits of the box represent the 75^th^ and 25^th^ percentile. The upper and lower vertical lines outside the box indicate maximum and minimum values in the data set. The unfilled circles outside the box denote potential outliers. Figure in inset depicts number of cattle in a grid as function of age. ^*^denotes statistically significant difference between male and female groups. ^NS^Not significant. Boxes bearing the identical superscripted letters are not statistically significantly different from each other. We have used these superscripted letters to show the difference only among the groups.

### Sex ratio

The estimated sex ratio was found to be 0.484:1 (male:female) based on 20 sampling grids. For each female there was less than one male in all studied 20 grids. The maximum (0.75:1) and the minimum (0.118:1) sex ratio were observed in Jai Stambh Area and WRS Colony, respectively (S2 Table).

### Age composition

We compared different age groups of the street cattle using Kruskal-Wallis test. Before employing this test we examined the normality (by Kolmogorov-Smirnov statistic) and homogeneity (by Levene statistic) of data. Results show that data were neither normally distributed (*p* < 0.05) nor homogenous (*p* < 0.05). This result did not fulfill the criteria of parametric test; therefore, we performed the Kruskal-Wallis test. Mean difference was set to be significant at a level of *p* ≤ 0.05. Results show that there is significant difference in the number of sighted cattle among different age groups (H_*2*_ = 76.954; *p* < 0.001). The number of street cattle was in the order of mature > immature > calve (Fig 3 inset). Further, to know the differences among different combination of groups we analyzed the data using Mann-Whitney U test. Here the mean difference was set to be significant at a level of *p* ≤ 0.017. This *p* value was calculated using the Bonferani adjustment. For this, initially the used significant level for overall mean differences (0.05) was divided by the number of levels compared (3 in this case). Subsequently, we had a new significant level, i.e., 0.017 (0.05/3). Results indicate that the number of cattle in the mature group was significantly higher as compared to that of the immature (U = 1773.00; *p* < 0.001) and calve groups (U = 797.50; *p* < 0.001). Frequency of cattle in immature group was also significantly higher than the frequency of cattle in calve group (U = 1753.50; *p* < 0.001) (Fig 3 inset).

For the frequency of six different cattle groups, data were not normally distributed (*p* < 0.05) except for the distribution of heifers (*p* > 0.05). Data were also found to be non-homogenous (*p* < 0.05). Result of Kruskal-Wallis test shows that there was significant difference (*p* <0.05) in the frequency of different age groups of cattle (H_*5*_ = 95.283; *p* < 0.001) (Fig 3). Further, to know the differences among different combination of groups, data were analyzed using Mann-Whitney U test. Significant level was set at *p* ≤ 0.008 (we found this value by Bonferani adjustment, i.e., 0.05/6). The frequency of cow in 1 km^2^ grid was significantly higher than all other cattle groups (cow *vs* bull: U = 379.50; *p* < 0.001; cow *vs* heifer: U = 327.00; *p* < 0.001; cow *vs* immature male: U = 191.00; *p* < 0.001; cow *vs* female calve: U = 97.50; *p* < 0.001; cow *vs* Male calve: U = 65.50; *p* < 0.001). Frequency of bull was significantly higher as compared to that of the immature male cattle (U = 500.00; *p* = 0.004), female calve (U = 355.50; *p* < 0.001), and male calve (U = 279.00; *p* < 0.001), but did not differ significantly with that of heifers (U = 755.00; *p* = 0.664). Further, frequency of heifer was significantly greater as compared to that of the immature male cattle (U = 508.00; *p* = 0.005), female calve (U = 339.50; *p* < 0.001), and male calve (U = 243.50; *p* < 0.001). The frequency of immature male cattle was significantly more as compared to that of the male calve (U = 527.50; *p* = 0.008), but did not differ significantly with that of female calve (U = 643.00; *p* = 0.127). The frequency of female and male calves did not differ significantly (U = 652.00; *p* = 0.148) (Fig 3).

## Discussion

In the urban areas, people share the public places with street cattle and street dogs. Number of cattle and dog is continuously increasing on the street/road/highways. There are several studies regarding free ranging urban dogs and their population management program in many urban areas of the world [19,22,33–36]. However, there is lack of information on the population of street cattle and on their ecological and behavioural aspects. We found only two publications while searching on the SCOPUS database [9,11]. Both studies highlighted circadian rhythms in various behavioural variables of street cattle. One of these studies also estimated the population of street cattle in 12 sampling sites in one of the Indian cities (Sambalpur), but did not defined the area of the studied sites [9].

The human population is 1.87 million in Raipur city [30] and we estimated population of street cattle is 34623. Based on the finding of this study, we can interpret that for every 54 people there is 1 street cattle in the city. This human:cattle ratio is an alarming and challenging situation especially for the Raipur Municipal Corporation. The number of street cattle is on the rise making the chances of man-cattle conflict higher. Every day, there are a large number of vehicular accidents attributed to the involvement of cattle on the street [15] and even cattle also get injured [37]. Unfortunately, we could not find any data regarding the human-cattle conflict on the road in Raipur city. The State Government and Municipal Corporation, Raipur have initiated work to find out ways to mitigate this cattle menace.

In this study, we observed and evaluated the density and population of street cattle using two different scientific survey protocols. Estimating population by using density of direct head count method in terms of animal per km distance is a novel method. This method has been recently used to estimate the density of street dogs in different countries in cohort studies [25,26]. The authors evaluated the density of street dog in terms of number of street dog per km distance. A recent study suggested that the total street dog population can be estimated by multiplying the density to the total length of road in particular area where population is to be estimated [25]. They also recommended for conducting comparable studies to validate this novel method.

Advanced PCRCM is a well established relevant method and has been successfully used in different studies to estimate the population of different species [16–19,21,22,36]. Most recently, it has been used to estimate the population of street cattle in 12 study sites of Sambalpur city, Odisha, India and the authors recorded about 146 street cattle on the 12 studied sites [9]. They did neither define the size of area nor estimate the total population of cattle in the city. In this study we used 20 randomly chosen 1 km^2^ sampling grids and computed the total estimated population of street cattle in total area of Raipur city.

In this study, we found that the estimated population of cattle on the street of Raipur city was about 11808 (by Line transect sampling) and 11198 (by Block count sampling) using direct head count method; and the estimated total population of street cattle in Raipur city was 35150 (by Line transect sampling) and 34623.20 (by Block count sampling) based on advanced Photographic Capture-Recapture Method (PCRCM) & Lincoln index formula (Table 4). The estimated population of direct head count method will always be lower as compared to the PCRCM. The possible reasons for lower estimation by direct head count are: (a) this method was based on average number of street cattle at two different days, (b) it does not include the cattle that could not be captured during photography or not visible within the observational window, (c) some cattle were not present on the third day during recapture and could not be photographed, and (d) we only count the cattle that were seen on the street/road/highway and within adjacent 10 meters on both sides during our observation. However, the PCRCM estimates the total population of street cattle found in the city irrespective of their presence or absence during two observational days.

We can state that the direct head count method only estimates the average number of heads visible on the street at a given time, whereas the PCRCM estimates the total population of street cattle found in the city irrespective of their presence or absence during two observational days. The direct head count method appears to be adequate and appropriate for this and possibly for other comparable studies. This method can be used as an effective technique to estimate average population of any species on particular line transect at a given time. However, if someone is interested to estimate the total population of any species then we recommend using of the Lincoln index formula.

The basic difference between two above methods can be easily understand by a model diagram. Calculated density and estimated cattle population are presented in Fig 4 (a-b) by direct head count method and Fig 4 (c-d) by PCRCM.

**Fig 4.**
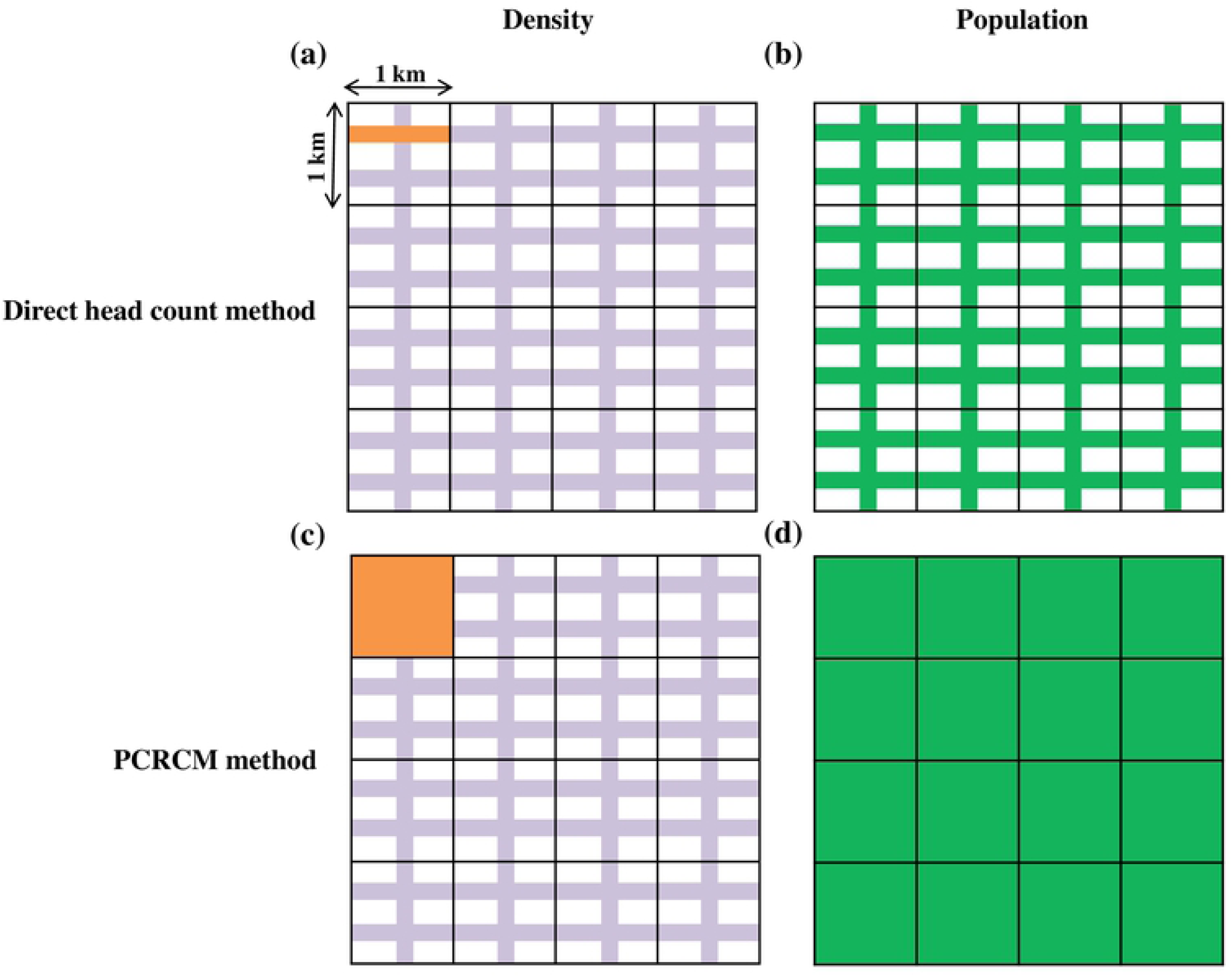
Model diagram - Density and population estimation of street cattle by two methods used in the study. (a) Density is represented (orange color) in terms of number of cattle per km distance and (b) estimated population in total road network (green color) by direct head count method. (c) Density (orange color) in terms of number of cattle per km^2^ area and (d) estimated population in total area (green color) by PCRCM. Road network is represented in light purple color.

Based on data emanated from our studies at 20 randomly chosen sampling grids, the average population density of street cattle in Raipur city was 7.28/km and varied from 0.40/km (min) to 23.58/km (max) (Table 2). The density depends on the extent of urbanization of the area. In our study, we found minimum density of street cattle in Jai Stambh Area. It is one of the most popular areas of the city having less number of residential household (3100) [28] and mainly used by people for occupational purposes. This area is mostly overcrowded between 0900 to 2300 h. The area is situated at the center of the city and has no open garbage disposal places. It may be one of the possible reasons for very less density of the street cattle in this area as there is very less opportunity for the cattle to find food there. On the other hand, the area Daganiya colony was having the maximum density of street cattle. This area is mostly occupied by people for residential purpose (7075 households) [28]. In this location cattle easily find food materials because this area has many open garbage disposal places. Cattle are mostly attracted towards the area as they find food materials easily.

Further, some earlier studies carried out in natural environments have reported male biased sex ratio in cattle population [38,39]. In contrast, we found female biased sex ratio of street cattle (male:female = 0.48:1). In the current study, we observed higher frequency of females not only in all studied grids (S2 Table), but also at different age levels (Fig 3).

Age composition analysis revealed that matured street cattle form 64.44% of the total population (S1 Fig). They are highly potential for breeding and they contribute significantly to the population growth of the cattle on the streets of the city. However, this hypothesis needs to be validated and confirmed through executing studies in future employing the same population estimation techniques.

## Conclusion

The estimated population of street cattle in Raipur city is 34623. There is a female biased sex ratio in the population of street cattle. Such a staggeringly high number of street cattle within the city are a challenge to the administrative authorities. Management of the free roaming cattle in the direction of making urban roads cattle free is beneficial to both the cattle and the citizens. The direct head count method that we adopted in this study is a novel technique and it always lower estimates the population as compare to the PCRCM. It can be useful for estimating the population of other animal species effectively.

## Data availability statement

All relevant data are within the manuscript and supporting information.

## Supporting information

**S1 Fig. Age composition (in percentage) in the population of street cattle in Raipur city (PDF)**.

**S1 Table. Estimated population of street cattle per km distance after applying the data based on line transect sampling into the Lincoln index formula at 20 different sampling grids in Raipur city (PDF)**.

**S2 Table. Sex ratio of street cattle in different studied grid: sex ratio is the ratio of males to females in a population (PDF)**.

## Acknowledgments

This work is a part of the Doctor of Philosophy thesis of one of the authors (BKS). He has been provided with fellowship by the University Grants Commission, New Delhi, India under the scheme of Joint CSIR-UGC NET for Junior Research Fellowship [Sr. No. 2121530765. Ref. No: 20/12/2015 (ii) EU –V dated 18/05/2016]. We are thankful to the Municipal Corporation of Raipur, India, for their cooperation and for providing us with the data on garbage bins installation and map of Raipur city. We record our especial thanks to Mr. Radhe Lal Markam for assisting us during the field study.

## Author contributions

**Conceptualization:** Bhupendra Kumar Sahu, Atanu Kumar Pati, Arti Parganiha

**Data curation:** Bhupendra Kumar Sahu

**Formal analysis:** Bhupendra Kumar Sahu, Arti Parganiha

**Investigation:** Bhupendra Kumar Sahu

**Methodology:** Bhupendra Kumar Sahu, Atanu Kumar Pati, Arti Parganiha

**Supervision:** Arti Parganiha, Atanu Kumar Pati

**Validation:** Arti Parganiha, Atanu Kumar Pati

**Visualization:** Bhupendra Kumar Sahu, Atanu Kumar Pati, Arti Parganiha

**Writing – original draft:** Bhupendra Kumar Sahu

**Writing – review & editing:** Bhupendra Kumar Sahu, Atanu Kumar Pati, Arti Parganiha

